## Supplementary Figures & Legends for "Physiological analysis of the mechanism of Ci transcription factor activation through multiple Fused phosphorylation sites in Hedgehog signal transduction"

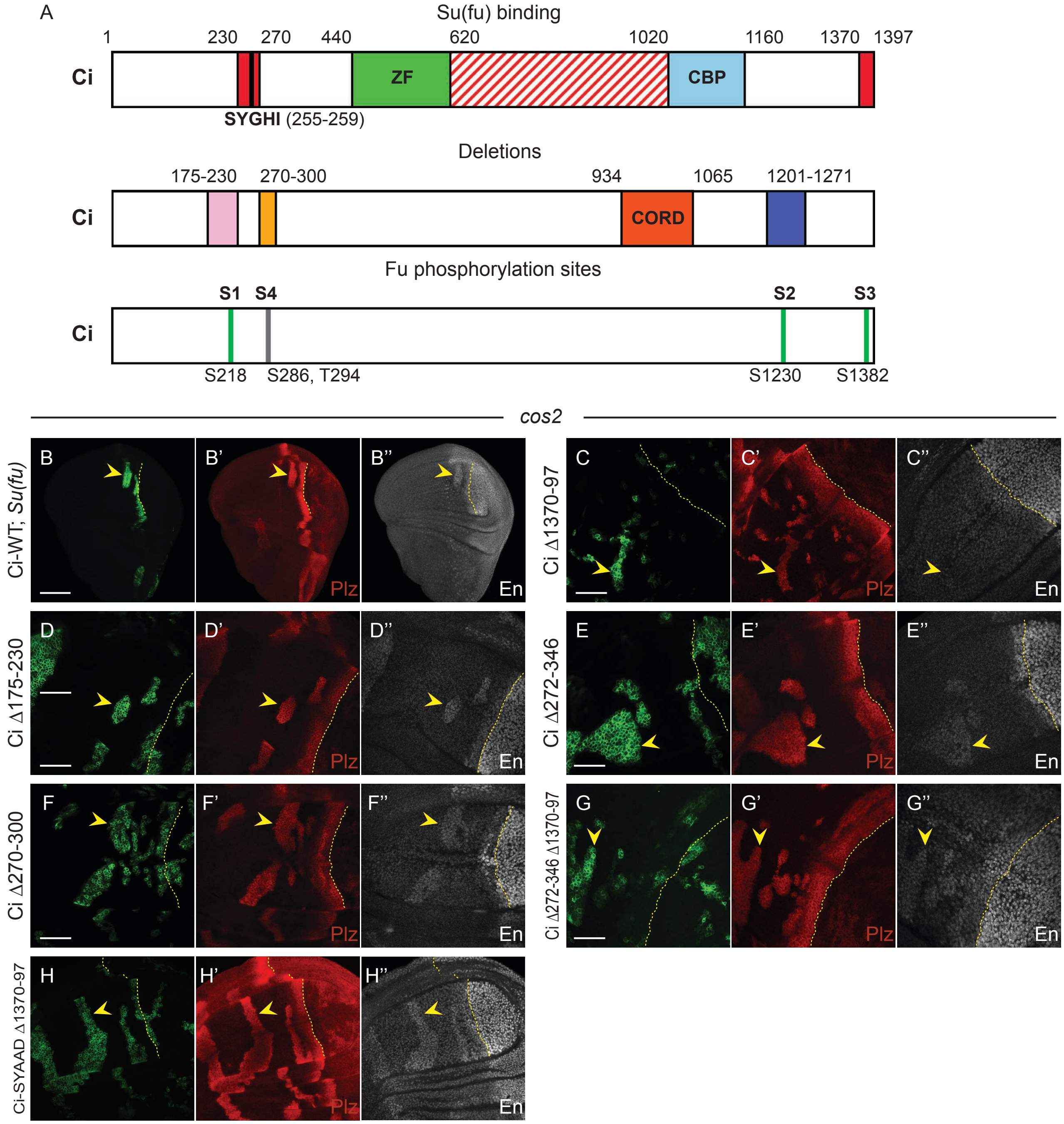


**Figure S1 (supplement to Fig. 1). Loss of Su(fu) binding sites or two SYGHI-adjacent regions increase Ci activity in *cos2* mutant clones: effects on En induction.**

(**A)** Key Ci features are illustrated. The top cartoon shows Su(fu) binding regions (red), the zinc finger domain (ZF, which binds DNA and can bind Cos2) and the binding region for CBP co-activator. The second cartoon shows deletions employed in this study (pink, yellow and blue) and the CORD Cos2 binding domain. The third cartoon shows Fu phosphorylation sites examined in this study. PKA sites

(S838, S856, S892) that promote Ci-155 processing, and a third Cos2 binding region (CDN; 346-440) are not shown. (**B-H**) Third instar wing discs (20x objective for (**B**) and 63x objective for all other images) with one copy of the indicated *ci* CRISPR allele, GFP marking homozygous *cos2* mutant clones (green; yellow arrowheads), and yellow dotted lines marking the AP border. (**B'-H'**) Ptc-lacZ expression (red) and (**B''-H''**) En protein (gray-scale) in the same discs. In (**B**) the entire wing disc lacks Su(fu) activity. Scale bars are 100μm for (**B**) and 40μm for all other images.

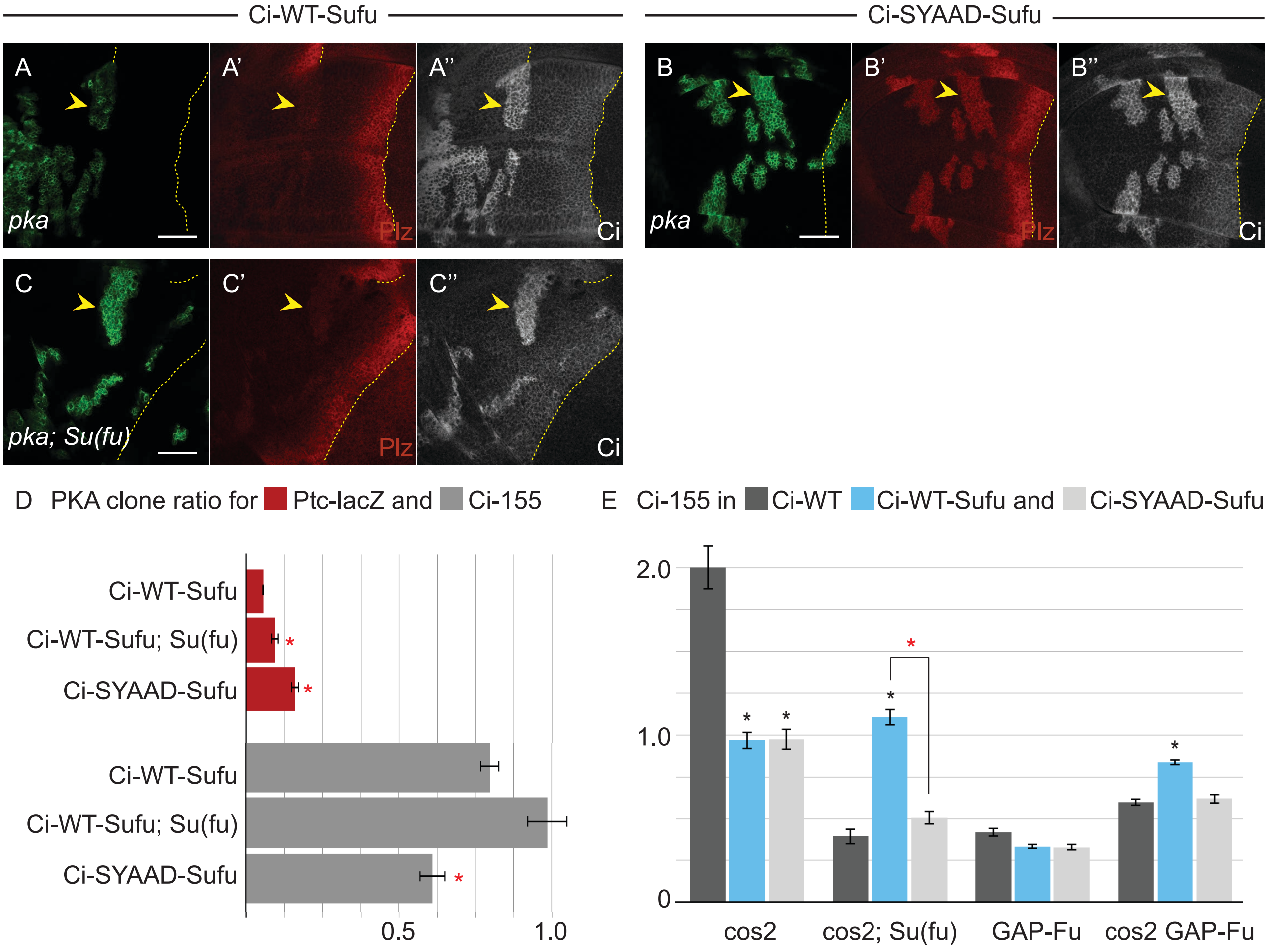


**Figure S2 (supplement to Fig. 4). Su(fu) inhibition is increased by covalent linkage to Ci but still requires non-covalent binding to the SYGHI region of Ci: effects in PKA mutant clones.**
(**A-C**) Third instar wing discs (63x objective) with one copy of (**A, C**) Ci-WT-Sufu or (**B**) Ci-SYAAD-Sufu, GFP marking *pka* mutant clones (green; yellow arrowheads), and yellow dotted lines marking the AP border. (**A'-C'**) Ptc-lacZ expression (red) and (**A"-C"**) Ci-155 expression (gray-scale) in the same discs. (**C**) Su(fu) activity was absent in the whole disc. Scale bars are 40μm. (**D**) Bar graph showing the average ratio of Ptc-lacZ intensity and Ci-155 intensity in clones relative to the AP border of wild-type control discs, together with SEMs (n values 94, 30, and 23, respectively, for each set of three genotypes). Differences with $p<0.005$ (Student's t test with Welch correction) are indicated for comparing to Ci-WT-Sufu in an otherwise wild-type disc (red asterisk). (**E**) Bar graph showing the average ratio of Ci-155 intensity in the indicated clones relative to the AP border of wild-type control discs, together with SEMs (n values 20, 5, 14, 10, 87, 19, 40, 54, 46, 47, 199, and 90); wing disc images and *ptc-lacZ* graph for these clones are in Fig. 4. Differences with $p<0.005$ (Student's t test with Welch correction) are indicated for comparing a Ci variant to Ci-WT (black asterisk) or for comparisons between bracketed pairs (red asterisk). Please see Materials and Methods for details of measurements and

expression of all experimental values relative to AP border values of control wild-type wing discs, and Fig4 plus S2_data supplementary information for raw data.

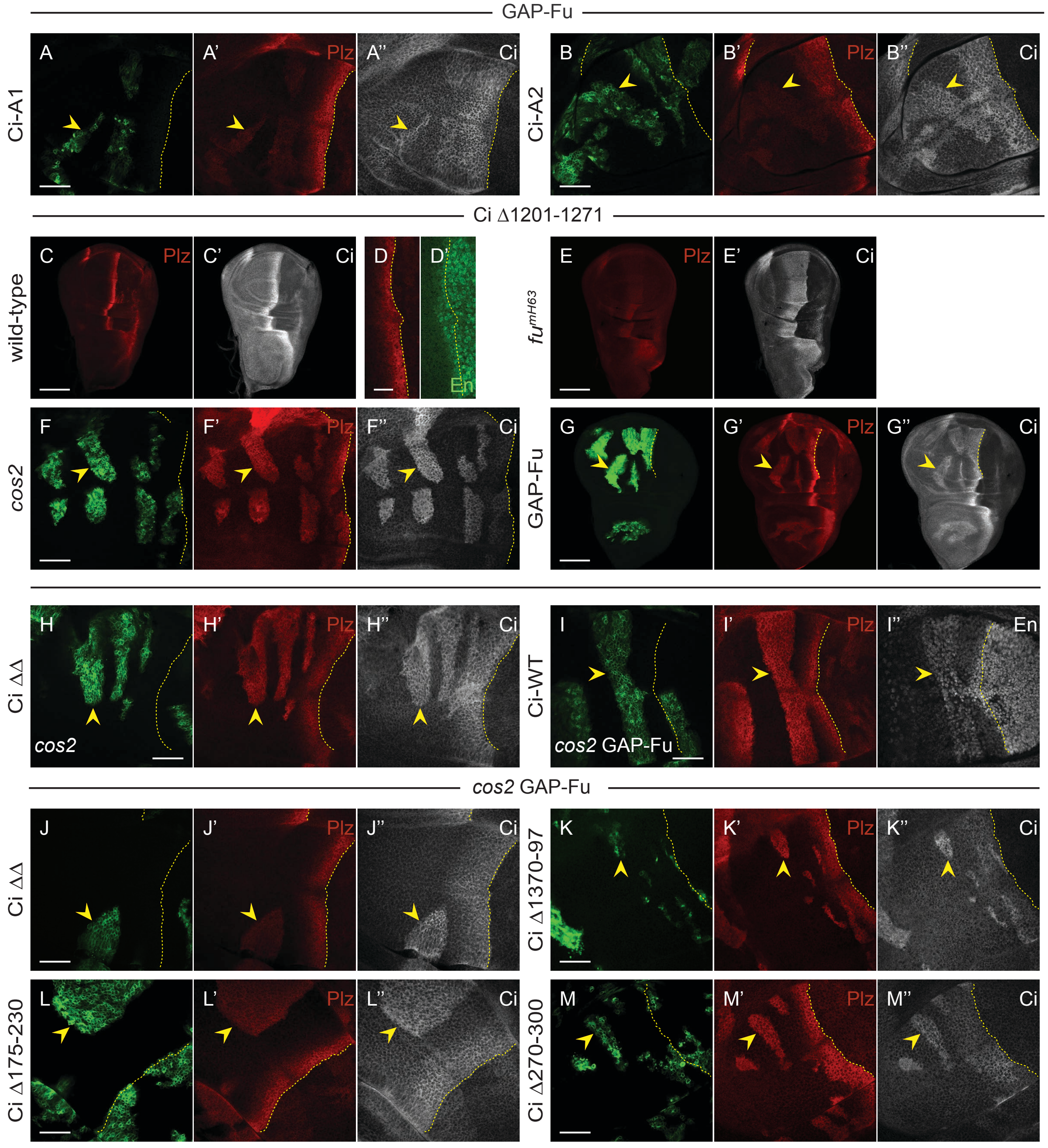


**Figure S3 (supplement to Figure 5). S218 and S1230 each contribute to activation by Fu and GAP-Fu reduces the activity of several Ci deletion variants in *cos2* mutant clones, as for Ci-A1A2.** (**A, B, F-M**) Third instar wing discs (20x objective for (**G**) and 63x objective for other images) with the named Ci variant, GFP marking the indicated clone types (green; yellow arrowheads), and yellow dotted lines marking the AP border. (**C-G**) all have Ci Δ1201-1271, while (**A, B**) have *GAP-Fu* clones and (**J-M**) have *cos2 GAP-Fu* clones. (**A‘, B‘, F‘-M‘**) Ptc-lacZ expression (red) and (**A“, B“, F“-H“, J“-M“**) Ci-155 expression (gray-scale) in the same discs. (**A, B, G, I-M**) *GAP-Fu* and *cos2 GAP-Fu* clones also lack *smo* activity. (**A, B**) Both S218A (Ci-A1) and S1230A (Ci-A2) reduced Ptc-lacZ induction by GAP-Fu. (**C-E**) Third instar wing discs with one copy of Ci Δ1201-1271, showing Ptc-lacZ (red) and (**C', E'**) Ci-155 expression (gray-scale) (20X objective) or (**D'**) En expression (green) (63X objective), resembling Ci-WT behavior, with the AP border marked by dotted yellow lines. (**F, G**) Induction of Ptc-lacZ and Ci-155 levels for Ci Δ1201-1271 also resembled Ci-WT in (**F**) *cos2* and (**G**) *GAP-Fu* clones. (**H-M**) The addition of GAP-Fu in *cos2* mutant clones (**I'**) increased Ptc-lacZ (compare to Fig. 1E) and (**I''**) En induction by Ci-WT but decreased Ptc-lacZ induction by (**H', J'**) Ci ΔΔ (which lacks residues 175-230 and 1201-1271) and by (**K'-M'**) Ci variants lacking residues 1370-97, 175-230 or 270-300 (compare to Fig. 1H, I, K). Scale bars are 40μm for (**A, B, F, H-M**), 100μm for (**C, E, G**) and 20 μm for (**D**).

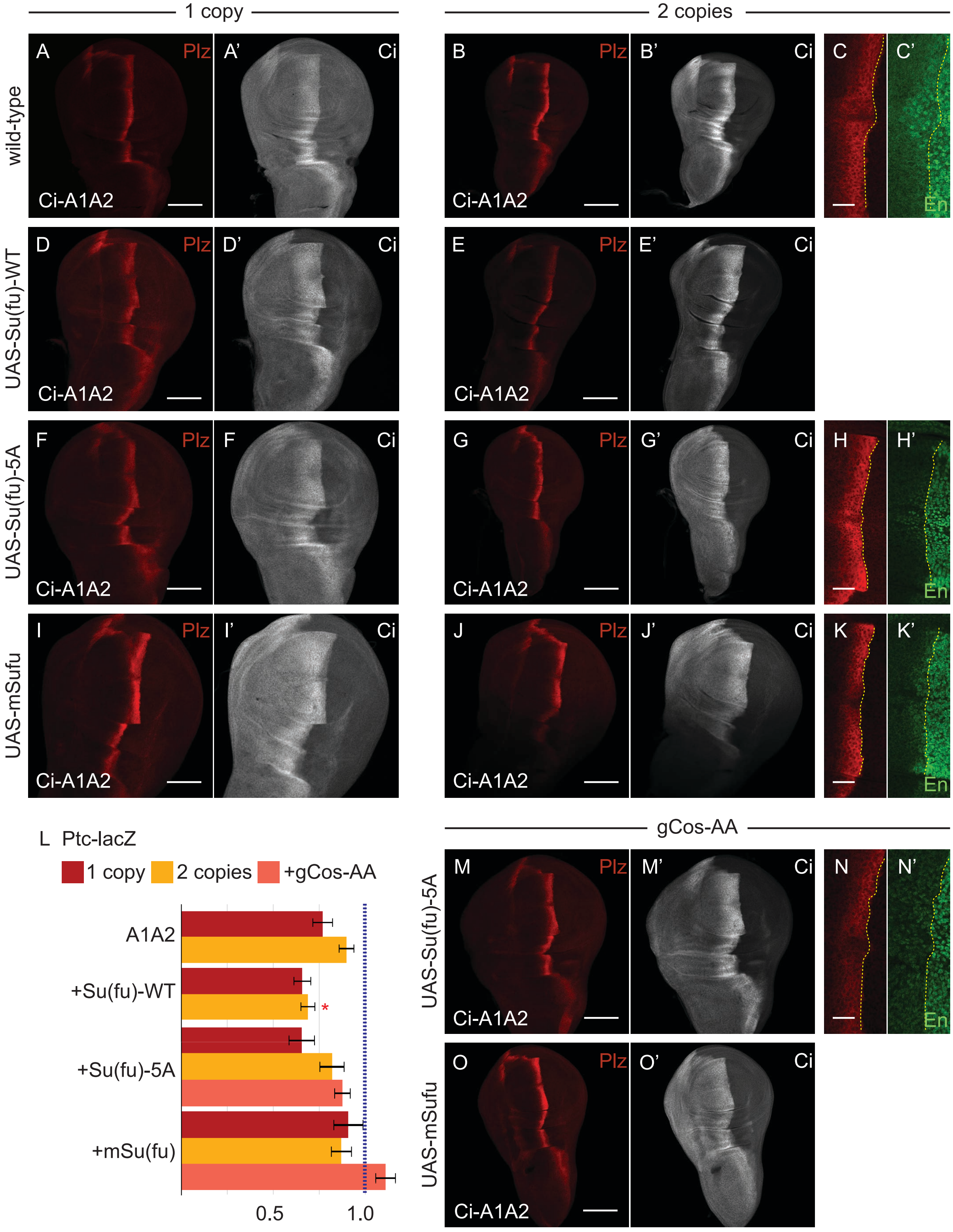


**Figure S4 (supplement to Figure 7). Loss of Fu sites in Su(fu) and Cos2 do not further reduce activity of Ci lacking S218 and S1230 Fu sites.**

(**A, D, F, I**) Third instar wing disc with one copy of Ci-A1A2 (*crCi-A1A2/ci*$^{94}$), showing (**A, D, F, I**) Ptc-lacZ (red) and (**A', D', F', I'**) Ci-155 expression (gray-scale) (20X objective). The discs are (**A**) otherwise wild-type or (**D, F, I**) *Su(fu)*$^{LP/LP}$ but expressing the indicated *UAS-Su(fu)* transgene using *C765-GAL4.* (**B, E, G,J, M-O**) Third instar wing disc with two copies of Ci-A1A2, showing (**B, E, G, J, M, O**) Ptc-lacZ (red) and (**B', E', G', J', M', O'**) Ci-155 expression (gray-scale) (20X objective) or (**C, H K, N**) Ptc-lacZ (red) and (**C', H', K', N'**) En expression (green) (63X objective; AP border marked by dotted yellow lines). The discs are (**B, C**) otherwise wild-type, (**E, G, J**) *Su(fu)*$^{LP/LP}$ but expressing the indicated *UAS-Su(fu)* transgene using *C765-GAL4* or (**M-O**) additionally lack endogenous *cos2* activity but contain one copy of the genomic transgene *gCos-AA* (encoding S572A S931A alterations). Su(fu)-5A has Fu site Ser residues substituted by Ala; mSufu encodes mouse Sufu. Scale bars are 20μm for (**C, H, K, N**) and 100 μm for all other images. (**L**) Bar graph showing the ratio of Ptc-lacZ intensity at the AP border of the named genotypes (with Cos-AA replacement of endogenous Cos2 in pink) relative the AP border of wild-type discs, together with SEMs (n= 29, 16, 27, 20, 17, 3, 22, 36, 28, 15 respectively for wing discs with Ci-A1A2). The blue dotted line at 1.0 marks the Ci-WT value. Differences with $p<0.005$ (Student's t test with Welch correction) are indicated for comparing *ptc-lacZ* at the AP border for Ci-A1A2 together with the indicated Su(fu) variant, or additional Cos2 variant, to Ci-A1A2 alone (separately for either one copy or two copies of Ci-A1A2) (red asterisk). Please see Materials and Methods for details of measurements and expression of all experimental values relative to AP border values of control wild-type wing discs, and FigS4_data supplementary information for raw data.

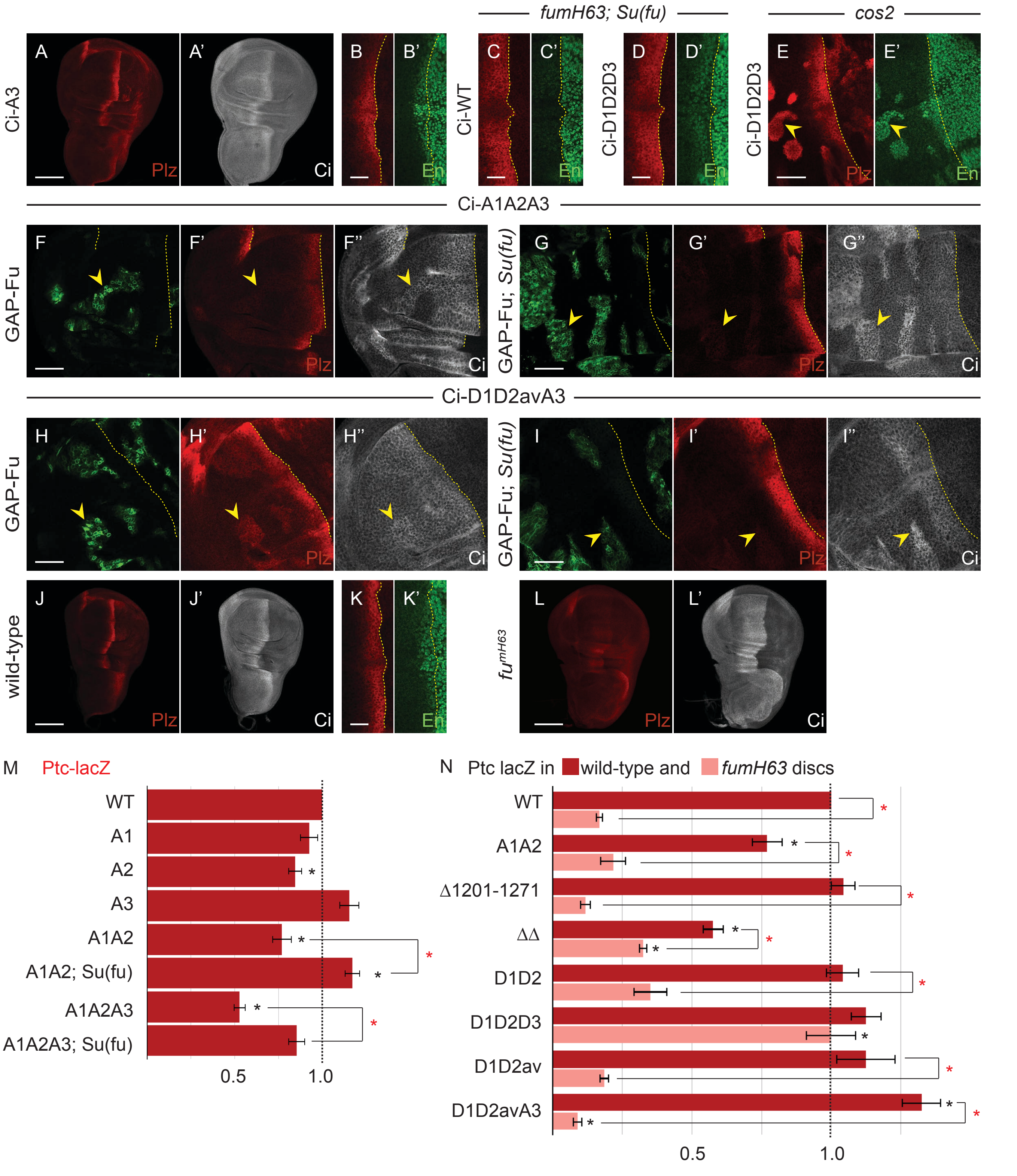


**Figure S5 (supplement to Figure 8). Contributions of S286, T294, S1382 and S1385 to Ci activation.**

(**A-D, J-L**) One copy of named Ci variants in (**A, B, J, K**) otherwise wild-type, (**C, D**) *fu*$^{mH63}$ *Su(fu)*$^{LP/LP}$ or (**L**) *fu*$^{mH63}$ third instar wing discs, showing (**A-D, J-L**) Ptc-lacZ (red), (**A', J', L'**) Ci-155 (gray-scale) or (**B'-**

**D', K'**) En expression (green, with the AP border marked by dotted yellow lines. (**E-I**) Third instar wing discs (63x objective) with one copy of (**E**) Ci-D1D2D3, with Ptc-lacZ (red) and En (green) expression shown in *cos2* clones (yellow arrowheads), or with (**F, G**) Ci-A1A2A3 or (**H, I**) Ci-D1D2avA3, GFP marking (**F-I**) *smo GAP-Fu* clones (green; yellow arrowheads) in otherwise (**F, H**) wild-type or (**G, I**) $Su(fu)^{LP/LP}$ discs, and yellow dotted lines marking the AP border. Scale bars are 100μm for (**A, J, L**), 20μm for (**B, C, D, K**) and 40μm for all other images. (**M**) Bar graph showing the ratio of Ptc-lacZ intensity at the AP border for the named Ci variants and Su(fu) genotypes relative to the AP border of control discs, together with SEMs (n= 20, 17, 15, 29, 22, 23, and 7, respectively). (**N**) Bar graph showing the ratio of Ptc-lacZ intensity at the AP border for the named Ci variants in otherwise wild-type (red) and $fu^{m63}$ wing discs (pink) relative to the AP border of control discs, together with SEMs (n= 29, 15, 24, 39, 13, 17, and 12, respectively for wild-type and 38, 5, 7, 21, 7, 20, 14, and 10, respectively, for $fu^{mH63}$). The difference between pairs of values shows the contribution of Fu kinase activity at the AP border. (**M-N**) Differences with p<0.005 (Student's t test with Welch correction) are indicated for comparing a Ci variant to Ci-WT (black asterisk) or comparisons between bracketed pairs (red asterisk). Please see Materials and Methods for details of measurements and expression of all experimental values relative to AP border values of control wild-type wing discs, and FigS5_data supplementary information for raw data.
