## Supplementary material for "Physiological analysis of the mechanism of Ci transcription factor activation through multiple Fused phosphorylation sites in Hedgehog signal transduction": Ci variant Table

| Name of the Ci Variant | Mutation Description | Original Amino Acid Sequence | Mutant Amino Acid Sequence | Original DNA sequence | Mutant DNA sequence |
|---|---|---|---|---|---|
| **Ci-SYAAD** | SYAAD | SYGHI | SYAAD | TCT TAC GGT CAT ATT | TCT TAC GCT GCT GAT |
| **CI Δ1370-97** | deletion of 1370-97 | N/A | N/A | N/A | N/A |
| **Ci Δ175-320** | deletion of 175-230 | N/A | N/A | N/A | N/A |
| **Ci Δ272-346** | deletion of 272-346 | N/A | N/A | N/A | N/A |
| **Ci Δ270-300** | deletion of 270-300 | N/A | N/A | N/A | N/A |
| **Ci Δ272-346 Δ1370-97** | deletion of 272-346 and 1370-97 | N/A | N/A | N/A | N/A |
| **Ci-SYAAD Δ1370-07** | SYAAD | SYGHI | SYAAD | TCT TAC GGT CAT ATT | TCT TAC GCT GCT GAT |
| **Ci-WT-Sufu** | tethering of Su(fu) cDNA sequence to the C-terminus of Ci | N/A | N/A | N/A | N/A |
| **Ci-SYAAD-Sufu** | SYAAD, tethering of Su(fu) cDNA sequence to the C-terminus of Ci | SYGHI | SYAAD | TCT TAC GGT CAT ATT | TCT TAC GCT GCT GAT |
| **Ci-A1** | S218A, S220A | LSSSP | LASAP | CTT TCG TCA TCG CCT | CTT GCG TCA GCG CCT |
| **Ci-A2** | S1230A, T1232A, S1233A | SSMTSL | SAMAAL | TCA TCG ATG ACT AGC TTG | TCA GCG ATG GCT GCC TTG |
| **Ci-A3** | S1382A, S1385A | SLTSL | ALTAL | TCC CTT ACT TCC TTA | GCC CTT ACT GCC TTA |
| **Ci-A1A2** | S218A, S220A, S1230A, T1232A, S1233A | see Ci-A1 and Ci-A2 | see Ci-A1 and Ci-A2 | see Ci-A1 and Ci-A2 | see Ci-A1 and Ci-A2 |
| **Ci-A1A2A3** | S218A, S220A, S1230A, T1232A, S1233A, S1382A, S1385A | see Ci-A1, Ci-A2, and Ci-A3 | see Ci-A1, Ci-A2, and Ci-A3 | see Ci-A1, Ci-A2, and Ci-A3 | see Ci-A1, Ci-A2, and Ci-A3 |
| **Ci ΔΔ** | deletion of 175-230 and 1201-1271 | N/A | N/A | N/A | N/A |
| **Ci Δ1201-1271** | deletion of 1201-1271 | N/A | N/A | N/A | N/A |
| **Ci-D1D2** | S218D, S220D | LSSSP | LDSDP | CTT TCG TCA TCG CCT | CTT GAC TCA GAC CCT |
| | S1230D, T1232D, S1233D | SSMTSL | SDMDDL | TCA TCG ATG ACT AGC TTG | TCA GAC ATG GAT GAC TTG |
| **Ci-D1D2D3** | S218D, S220D, S1230D, T1232D, S1233D | see Ci-D1D2 | see Ci-D1D2 | see Ci-D1D2 | see Ci-D1D2 |
| | S1382D, S1385 | SLTSL | DLTDL | TCC CTT ACT TCC TTA | GAC CTT ACT GAC TTA |
| **Ci-D1D2AV** | S218D, S220D, S1230D, T1232D, S1233D | See Ci-D1D2 | See Ci-D1D2 | See Ci-D1D2 | See Ci-D1D2 |
| | S286A, T294V | ASAGLLNPMTP | AAAGLLNPMVP | GCA AGT GCA GGT CTG TTA AAT CCG ATG ACA CCA | GCA GCT GCA GGT CTG TTA AAT CCG ATG GTA CCA |
| **Ci-D1D2AVA3** | S218D, S220D, S1230D, T1232D, S1233D + S286A, T294V, S1382A, S1385A | See Ci-D1D2AV and Ci-A3 | See Ci-D1D2AV and Ci-A3 | See Ci-D1D2AV and Ci-A3 | See Ci-D1D2AV and Ci-A3 |
